# CryoEM Reveals Oligomeric Isomers of a Multienzyme Complex and Assembly Mechanics

**DOI:** 10.1101/2022.11.18.517149

**Authors:** Jane K.J. Lee, Yun-Tao Liu, Jason J. Hu, Inna Aphasizheva, Ruslan Aphasizhev, Z. Hong Zhou

## Abstract

Propionyl-CoA carboxylase (PCC) is a multienzyme complex consisting of up to six α-subunits and six ß-subunits. Belonging to a metabolic pathway converging on the citric acid cycle, it is present in most forms of life and irregularities in its assembly lead to serious illness in humans, known as propionic acidemia. Here, we report the cryogenic electron microscopy (cryoEM) structures and assembly of different oligomeric isomers of endogenous PCC from the parasitic protozoan *Leishmania tarentolae* (LtPCC). These structures and their statistical distribution reveal the mechanics of PCC assembly and disassembly at equilibrium. We show that, in solution, endogenous LtPCC β-subunits form stable homohexamers, to which different numbers of α-subunits attach. Sorting LtPCC particles into seven classes (i.e., oligomeric formulas α_0_β_6_, α_1_β_6_, α_2_β_6_, α_3_β_6_, α_4_β_6_, α_5_β_6_, α_6_β_6_) enables formulation of a model for PCC assembly. Our results suggest how multimerization regulates PCC enzymatic activity and showcase the utility of cryoEM in revealing the statistical mechanics of reaction pathways.

## Introduction

Multienzyme complexes are stable assemblies of multiple subunits of enzymes. These complexes are widespread [1–3], often involved in various metabolic pathways. One such multienzyme is propionyl-CoA carboxylase (PCC), which catalyzes the carboxylation of propionyl-CoA to form methylmalonyl-CoA, a precursor to the citric acid cycle intermediate succinyl-CoA [4]. As citric acid cycle is essential to cellular metabolism, PCC is found in bacteria [5], archaea [6], protozoa [7], plants [8], and animals [5]. In humans, inherited mutations in the genes encoding PCC may interfere with multienzyme assembly leading to catalytic dysfunction [9] and the metabolic disorder known as propionic acidemia [10]. Symptoms of propionic acidemia include metabolic acidosis, hyperammonemia, hypoglycemia, lethargy, vomiting, seizures, and possibly death [11,12]. Reflective of deep evolutionary conservation, this enzyme displays high protein sequence homology, exemplified by the sequence similarity between the protozoan *Leishmania tarentolae* PCC (LtPCC) and *Homo sapiens* PCC (HsPCC) (Figure S1).

Prior structural studies have resolved a 3.2 □Å crystal structure of a recombinant PCC chimera (PCC^chi^), where its α-subunit is from *Ruegeria pomeroyi* and β-subunit is from *Roseobacter denitrificans* [5]. A cryoEM structure of the recombinant HsPCC has been determined at 15 Å resolution showing similar architecture to PCC^chi^ [5]. More recently, a 3.48□Å cryoEM structure of the recombinant *Methylorubrum extorquens* PCC (MePCC) was used to guide the design of a new-to-nature enzyme for improved CO_2_ fixation [13]. In the aforementioned cryoEM studies, PCC was found to oligomerize as an α_6_β_6_ dodecamer.

For PCC to catalyze carboxylation, the enzyme must first be biotinylated [14]. Following biotinylation, the multienzyme acts in two steps involving α- and β-subunits. In the first step, biotin is carboxylated at an α-subunit active site with bicarbonate as the carbon dioxide donor upon concomitant ATP hydrolysis [5,15]. In the second step, the carboxylated biotin is translocated to the corresponding β-subunit active site, and the carboxyl group is transferred from biotin to form methylmalonyl-CoA [5,15].

Given that assembly and disassembly propensities of multienzyme complexes may influence catalytic efficiency, it is crucial to model their assembly mechanisms. CryoEM enables the observation of multienzymes in different assemblies and to reconstruct different oligomeric isomers. Unlike X-ray crystallography, endogenous proteins in different stages of assembly can be classified and counted in cryoEM micrographs to reveal the statistical mechanics of chemical reactions, along with obtaining high-resolution atomic structures.

Here, we report three structures of endogenous LtPCC: an α_6_β_6_ dodecamer, α_5_β_6_ undecamer, and α_4_β_6_ decamer. We utilize the α_5_β_6_ and α_4_β_6_ architectures of PCC to demonstrate that PCC oligomeric isomers differ only in their number of α-subunits. We devised a sorting method to calculate the number of LtPCC proteins with the same oligomeric formula, from α_0_β_6_ to α_6_β_6_. From this statistical information, we characterized the dynamics of PCC’s molecular assembly system and reaction mechanics in solution.

## Results

### LtPCC α_6_β_6_ dodecamer and domain organization

To capture cryoEM structures of various biotin-binding complexes, we performed streptavidin pull-down of endogenous complexes from *L. tarentolae* mitochondrial lysate and determined their structures. One subset of these structures has a three-fold symmetric architecture reminiscent of carboxylases [5,15–17] and, using the cryoID approach [18], we confirmed its identity as propionyl-CoA carboxylase from *L. tarentolae* (LtPCC).

With D3 symmetry, we obtained a map of the LtPCC α_6_β_6_ dodecamer at 3.2 Å resolution. As a dodecamer, PCC contains two layers of α-subunits sandwiching a β-homohexamer. Each layer of α-subunits contains three monomeric α-subunits while the six β-subunits in the β-homohexamer are arranged into a two-layered cylinder (Figures 1A-B). Thus, the architecture of α_6_β_6_ LtPCC is the same as that of PCC^chi^, MePCC, and HsPCC [5,13].

**Figure 1.**
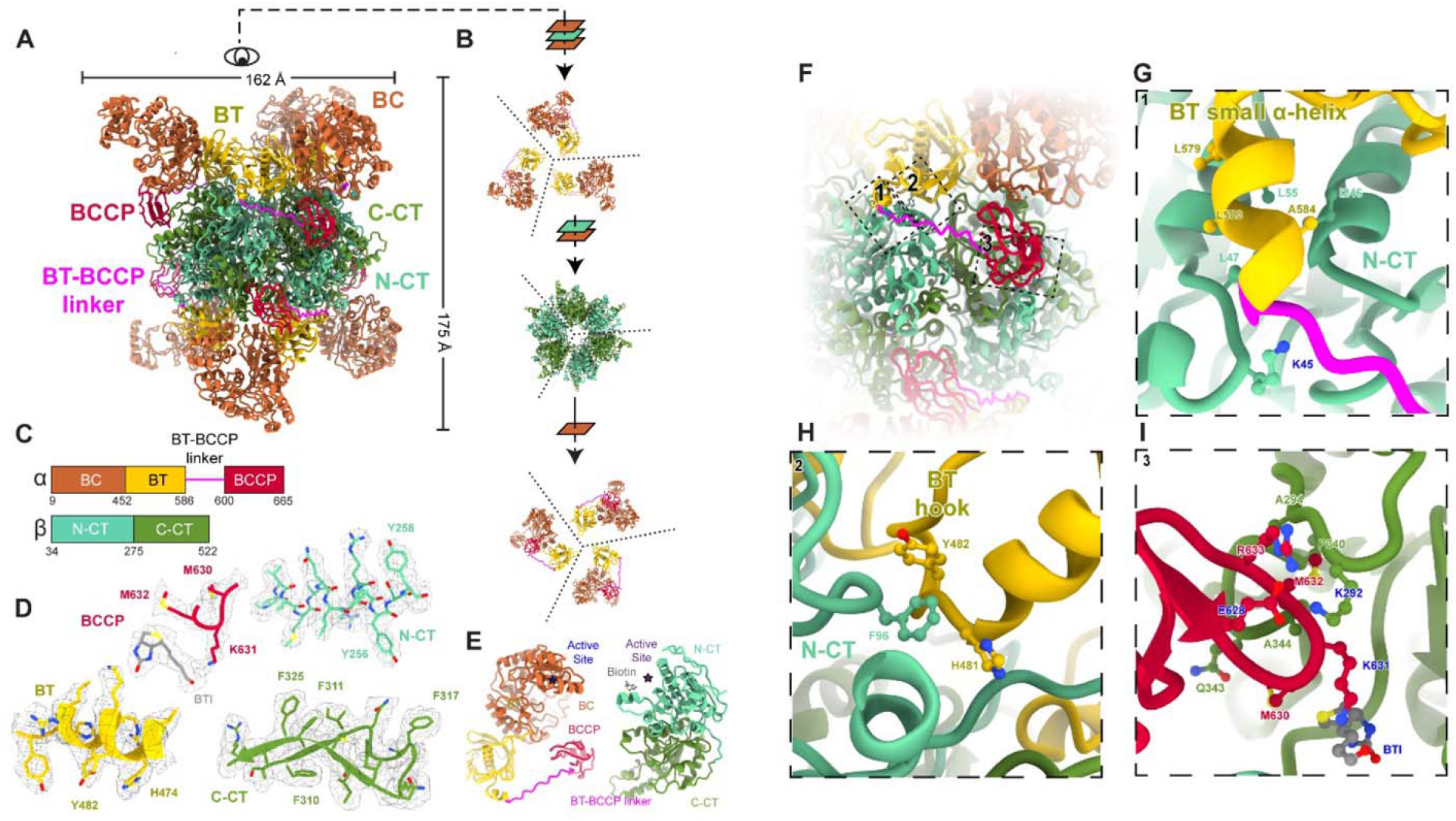
Structure of the LtPCC α_6_β_6_ dodecamer and the α-β binding site. (A) Atomic model of LtPCC α_6_β_6_ dodecamer, shown as ribbons and colored by domains. (B) A bird’s eye view of the atomic model in (A). Dashed lines indicate the boundaries between neighboring subunits. The β-homohexamer contains two-layers of subunits. (C) Domain organization of LtPCC α- and β-subunits; the color scheme is used throughout the manuscript. (D) Representative cryoEM densities superimposed with the atomic model of LtPCC shown as ribbons and sticks. (E) Atomic model of a α-subunit and a β-subunit of LtPCC colored by domains and shown as ribbons, with active sites labeled. Residues at the α-β binding site are boxed with dotted lines in (F) and labeled in zoomed-in views (G-I). Residues that form hydrogen bonds are labeled in blue. BTI stands for biotin.

The α-subunit of LtPCC contains three domains: a biotin carboxylase (BC) domain, BC-CT mediating (BT) domain, and biotin carboxyl carrier protein (BCCP) domain. A BT-BCCP linker connects the BT and BCCP domains, allowing BCCP to transport biotin between the α- and β-subunits. As in PCC^chi^, the active sites in LtPCC are positioned 55 Å apart; therefore, the BCCP domain must translocate between the active sites, as proposed by the swinging-domain model [5,15]. The β-subunit is composed of the structurally homologous N-carboxyltransferase (N-CT) and C-carboxyltransferase (C-CT) domains [5,13] (Figures 1C-E).

In carboxylases, biotin attaches to a conserved lysine residue in the alanine-methionine-lysine-methionine (AMKM) biotinylation motif within the BCCP domain [19,20]. Accordingly, we observed clear density for biotin next to Lysine 631 of the AMKM motif in the β-subunit active site. The biotin’s carboxyl group, which is the point of covalent attachment to lysine, neighbors the lysine residue. There is weak density connecting lysine and biotin (Figure 1D), suggesting that some LtPCC assemblies are covalently biotinylated. CoA was not observed at its β-subunit binding site.

### α-β binding sites within LtPCC

Using our 3.2 Å map of the dodecamer, we analyzed inter-subunit interactions within LtPCC. Interactions between LtPCC α- and β-subunits enable complex assembly (Figures 1F-I) and make up the α-β binding site, with a binding affinity of −12.8 kcal mol^-1^. No interactions exist among LtPCC α-subunits. There are seven hydrogen bonds between α- and β-subunits. Three hydrogen bonds occur between the BCCP domain of the α-subunit and the β-subunit, and two hydrogen bonds form between the BT-BCCP linker and the β-subunit. Glutamate 628 from the BCCP domain forms two hydrogen bonds with Lysine 292 from the β-subunit (Figure 1I). There is another mainchain-mainchain hydrogen bond between the mainchain of Arginine 633 in the BCCP domain and the β-subunit. Glutamate 628, the biotin-attachment residue Lysine 631, and Arginine 633 form a U-shape with Lysine 631 in the middle, with their three hydrogen bonds stabilizing the U-shaped loop to hold the lysine residue in proximity to biotin (Figure 1I). Lysine 45 from the β-subunit forms two hydrogen bonds with the BT-BCCP linker mainchain (Figure 1G), stabilizing the linker. These interactions hold the BCCP domain in place near the β-subunit.

Among the α-subunits and β-subunits, residues contributing to the α-β binding site primarily occur along the small α-helix of the BT domain with the N-CT domain, the BCCP domain with the C-CT domain, and the BT domain hook with the N-CT domain (Figures 1F-I). The small α-helix of the BT domain, consisting of residues 578-585, interacts with the N-CT domain through (iso)leucine-mediated hydrophobic interactions (Figure 1G). A loop (from residues 628–633) of the BCCP domain contacts two C-CT domain loops (one from residues 340–344 and the other from 292–294) (Figure 1I). The α-subunit BT hook, formed by residues 481–491, interacts with the N-CT domain of its neighboring β-subunit via the π-π stacking among the three aromatic rings of Tyrosine 482, Phenylalanine 96, and Histidine 481 (Figure 1H). Taken together, hydrophobic interactions and hydrogen bonds contribute to the α-β binding site to stabilize the α-subunits onto the β-homohexamer.

### LtPCC α_4_β_6_ and α_5_β_6_ oligomeric isomers differ from α_6_β_6_ dodecamer in α-subunit occupancies

In the α_6_β_6_ dodecameric structure, α-subunits disappear when observed at high density threshold, indicating that α-subunits are either flexible and/or have lower occupancy (Figure S2). The 2D class averages also contain classes with missing α-subunits (Figure S3). To sort out LtPCCs with missing α-subunits, we performed symmetry-relaxed 3D classification and determined two additional LtPCC structures (Figures 2A-C): α_5_β_6_ undecamer, α_4_β_6_ decamer. The α_5_β_6_ undecamer lacks one α-subunit, and the α_4_β_6_ decamer lacks two subunits on opposite sides of the LtPCC complex. In our decamer, undecamer, and dodecamer, the conformation of individual subunits is preserved. In other words, the structures of these complexes differ from one another only in the number and positioning of α-subunits. It follows that a single α-β binding site gives rise to a binary choice of the binding site being occupied or unoccupied (Figures 2A-C).

**Figure 2.**
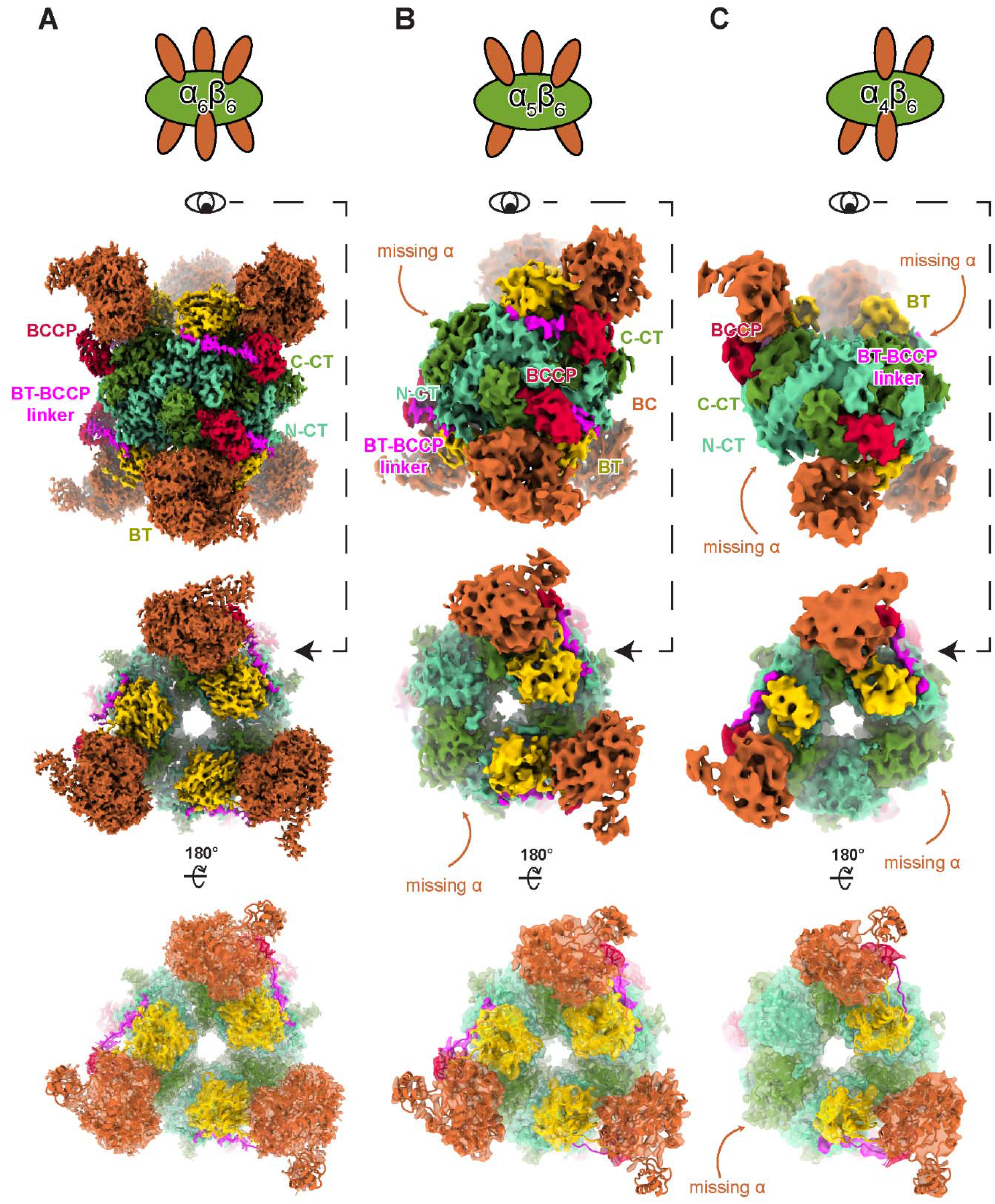
LtPCC conformations differ only in occupancies of the α-β binding sites. Cartoon representations of α_6_β_6_ (A) α_5_β_6_ (B) and α_4_β_6_ (C) cryoEM densities, with α-subunits in orange and β-subunits in green (top row). Two views of the cryoEM densities for α_6_β_6_ (A) α_5_β_6_ (B) and α_4_β_6_ (C) colored by domain as in Figure 1C (middle rows), superimposed with their respective atomic models represented as ribbons (bottom row).

### Reaction mechanics and model of a multienzyme assembly and disassembly

To facilitate the description of the various structures we observed, we introduce three terms: oligomeric formula, oligomeric isomer, and structural conformation. An oligomeric formula is akin to a chemical formula, and a LtPCC oligomeric formula can be written as α_n_β_m_. Multiple oligomeric isomers could share the same oligomeric formula but have different arrangements of subunits in space. Though the architectural arrangement of subunits in an oligomeric isomer is unique, each subunit could assume different structural conformations. In the case of LtPCC, as described above, each oligomeric isomer has one, and only one structural conformation.

The above-mentioned three structures are only three possible isomers that LtPCC could have in solution. There are sixteen possible different LtPCC oligomeric isomers represented by seven oligomeric formulas (one isomer for α_6_β_6_, α_5_β_6_, α_1_β_6_ and α_0_β_6_, and four for α_4_β_6_, α_3_β_6_ and α_2_β_6_). Even though there is only one structural conformation for each oligomeric isomer, the large number of potential oligomeric isomers and the structural similarity among these oligomeric isomers present a technical challenge in relying on traditional classification methods to sort out all these isomers.

To tackle this problem, we used a sorting method to determine the number of particles belonging to each oligomeric formula by classifying its α-subunits. As indicated above, for each oligomeric isomer, the α-β binding site is either occupied or unoccupied. We searched for the presence or absence of an α-subunit at each of the six α-β binding sites on a PCC particle, giving rise to 2^6^ possible permutations, sharing seven oligomeric formulas and sixteen oligomeric isomers (Figures 3A-B). We counted the number of occupied α-β binding sites in each PCC particle (Table 1), to assign each particle to its corresponding oligomeric formula.

**Figure 3.**
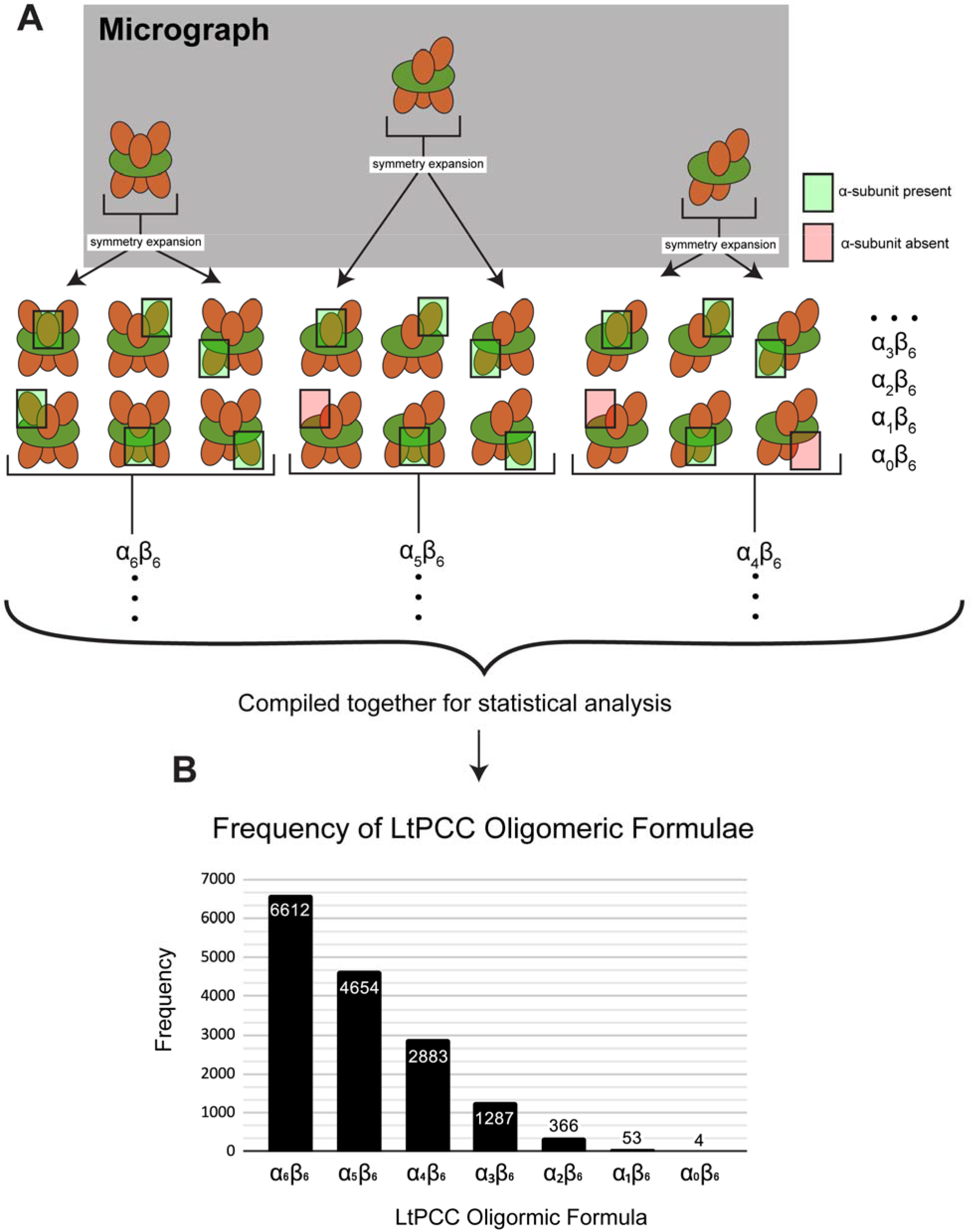
Sorting method to determine LtPCC oligomeric formula distributions. (A) Cartoon representation of the sorting method, with α-subunits in orange and β-subunits in green. (B) Frequency graph of LtPCC oligomeric formulae calculated from the sorting method.

The sorting method enables calculation of the frequency distribution of PCC particles belonging to different oligomeric formulae. We found that most endogenous LtPCCs exist as α_6_β_6_, and in descending frequency, LtPCC also exhibit oligomeric isoforms with the oligomeric formulas α_5_β_6_, α_4_β_6_, α_3_β_6_, α_2_β_6_, α_1_β_6_, and α_0_β_6_ (Figure 3B). Previous crystal and cryoEM structures of PCCs only captured its α_6_β_6_ dodecamer, which does not account for all PCC structural conformations. The presence of PCC oligomeric isomers is corroborated by previous biochemical studies that suggest PCCs might assemble as tetramers [21,22]. Given that we only observe β-homohexamers and α-β complexes, the assembly of functional PCCs likely occurs after the assembly of β-homohexamers (Figure 4).

**Figure 4.**
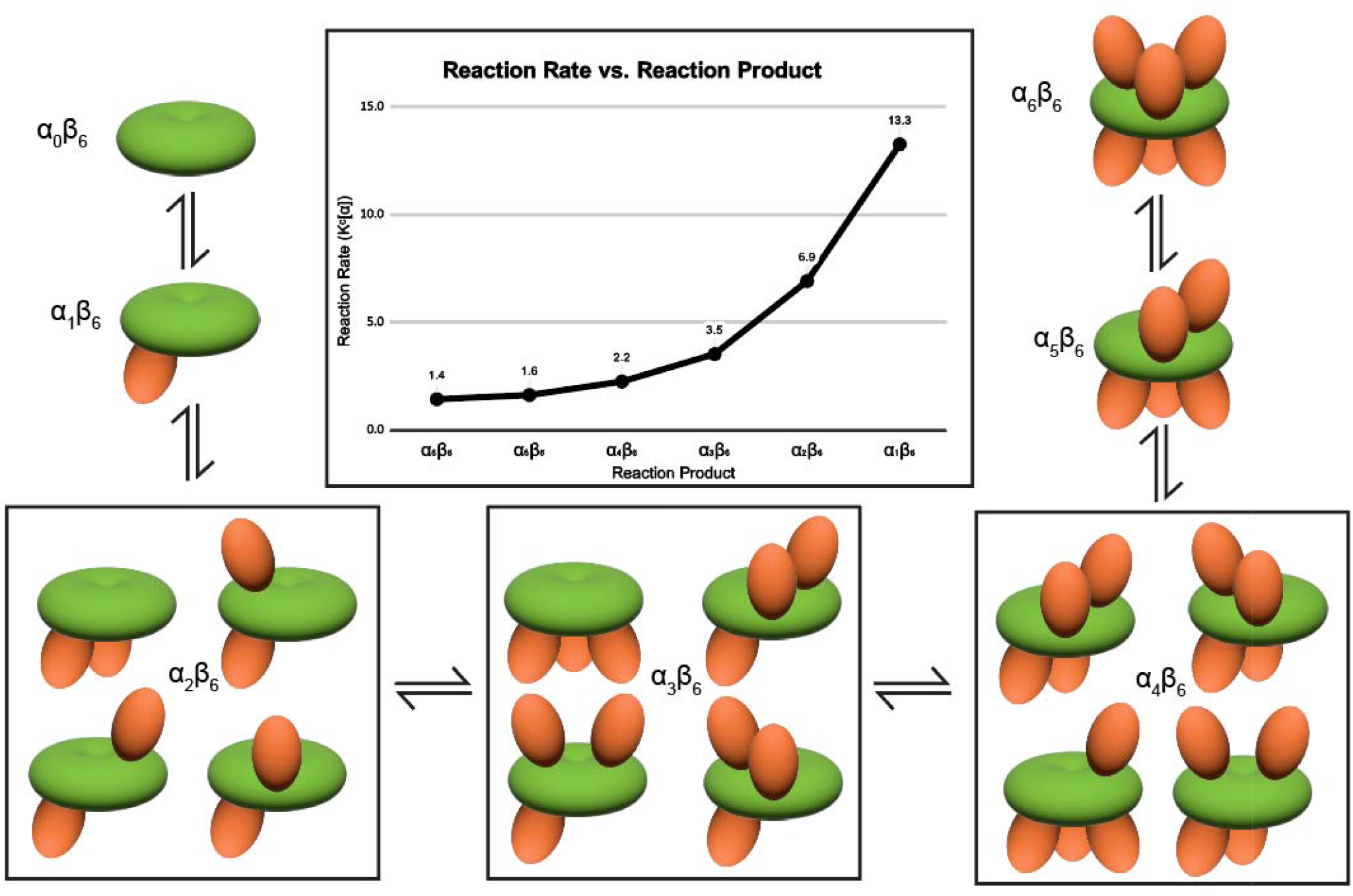
Reaction mechanics of LtPCC. In the center is the graph of LtPCC reaction rate versus reaction product, showing exponential decay of reaction rate with increasing numbers of α-subunits in the reaction product. Surrounding the graph is the reaction diagram of LPCC assembly/disassembly, with oligomeric isomers of each oligomeric formula grouped together. α-subunits are colored in orange and β-subunits are colored in green.

The solution in our sample is at an equilibrium state, as it has mixed and settled for hours before freezing. The equilibrium constant *K_c_* can be obtained by calculating the ratio of product concentration to reactant concentration. The assembly of LtPCC can be represented by this reaction:

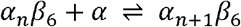

The equilibrium constant *K_c_* can be found by:

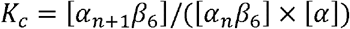

*α* and *β* represent α- and β-subunits, respectively; square brackets indicate concentration; *n* is an integer that denotes the number of α-subunits in LtPCC. We are unable to measure the concentration of α-subunits because single α-subunits are small and flexible, and therefore cannot yet be recognized in cryoEM images. Nevertheless, we can calculate the product of [*α*] and *K_c_* as an attribute of the PCC assembly reaction. We assume that [*α*] remains constant throughout due to the assembly and disassembly of α-subunits at equilibrium (Figure 4).

The plot of *K_c_*[*α*] as a function of *n* (Figure 4) shows that *K_c_*[*α*] decreases exponentially with increasing *n*. With [*α*] remaining constant, this means that particles with a smaller *n* value have greater tendency to attach new α-subunits compared to particles with a larger *n* value. This can be partially explained by complexes with more α-subunits having less reaction sites for α-subunit attachment.

## Discussion

Our present work shows promise of utilizing cryoEM for statistical analysis of thermodynamics and structural dynamics to understand the behaviors and assembly of biological complexes. By simultaneously determining different structural conformations of a protein, we can discern its reaction mechanics and calculate the rate of conversion between its oligomeric isomers. The future prospect of experimentally determining statistical mechanics presents exciting opportunities for a deeper understanding of the catalytic mechanisms of multienzyme complexes and the working-cycle of molecular machines.

Crystallography studies have elucidated β-homohexamer structures with no α-subunits in various carboxylases, including PCC [16,23]. Therefore, we propose that β-homohexamers are stable in solution, and the lack of interactions between PCC α-subunits allow for a continuous assembly and disassembly of α-subunits from β-homohexamers. As a dodecamer, PCC contains six pairs of active sites per enzyme available for catalysis. In other oligomeric isomers, at least one active site cannot participate in catalysis, and so all non-dodecameric PCCs are less catalytically active than the dodecamer. As the concentration of α-subunits increases, to maintain equilibrium, the reaction will favor the formation of PCC isomers with more α-subunits. The maintenance of β-homohexamers allows for quick regulation of PCC activity, where only α-subunits need to assemble.

Isomers arise from protein conformational heterogeneity, and in other systems, an oligomeric isomer can have multiple structural conformations. In theory, cryoEM allows one to determine all these structural conformations and derive their reaction kinetics. An example of a dynamic macromolecular machine is the spliceosome. During transcription, the spliceosome splices introns from pre-messenger RNAs. Throughout the splicing cycle, some spliceosome components are displaced while others assemble, and assembled components undergo conformational changes. Though structures of the spliceosome at different stages of the splicing cycle exist, the assembly kinetics of this process remains to be resolved [24–28]. Similarly, ribosomes undergo dynamic compositional and conformational changes during translation; despite structures of multiple ribosome states, information about its reaction kinetics remains unknown [29]. The complexity of resolving all isomers in multi-subunit systems would require imaging and computational resources that are financially prohibitive at present. The relatively simple system with just two molecules each of a single conformation already has sixteen oligomeric isomers. The presence of D3 symmetry further allows application of a sorting method to count the numbers of particles sharing the same oligomeric formula without having to solve all structures exhaustively using an enormous cryoEM dataset.

Currently, the difficulty in seeing small molecules (<50 kDa) with cryoEM makes it unfeasible to directly find the rate constant of molecular assembly involving small components, though future developments should enable the determination of such components by cryoEM [30,31]. Just like how Google’s AlphaFold [32] came about in 2020 to solve the previously computationally-prohibitive protein-folding-prediction problem, future cryoEM imaging and computational resources should enable determination of all structural conformations in a complex assembly to derive reaction kinetics. In fact, if the cells are thin enough, we not only can determine structural conformations in solution but also in situ [29,33,34]. Such prospects offer exciting opportunities for experimentally “visualizing” statistical mechanics within an enormous conformational space, and to assist drug design in targeting the rate-limiting step of a complex of interest’s assembly [35–38]. Taking advantage of the limited number of conformational isomers of PCC, the current work showcases the utility of cryoEM beyond determining static structures towards statistical analysis of thermodynamics and structural dynamics.

## Materials and Methods

All methodology except cryoEM image processing is the same as here [39].

### Lead contact

All information and requests for further reagents and resources should be made to and will be fulfilled by the lead contact, Z. Hong Zhou.

### Experimental model and subject details

We grew *L. tarentolae* cells in brain heart infusion media at 27°C. The media was supplemented with 5 mg/L of hemin. We harvested the cells at ~2×10^8^ cell/ml during the late-exponential growth phase.

### Preparation of mitochondrial lysate

We enriched mitochondrial fraction through hypotonic cell lysis, and used RenoCal76 density gradients for the sequential separation of membrane-containing fractions [40]. By sonication at 24W for 15 seconds and centrifugation at 30,000 RPM in a SW55 rotor for 15 minutes, we lysed mitochondrial pellets in 1 ml of pH 7.3, 50 mM HEPES, 150 mM KCl, 2 mM EDTA, 1% NP40, and 50 μL of 20x complete protease cocktail. We recovered and separated the supernatant on a continuous 10-30% gradient glycerol in pH 7.3, with 20 mM HEPES, 100 mM KCl, and 1 mM EDTA, prepared at 72,000 g for 15 hours, in SW28/32 Setton clear tubes. We collected glycerol gradient fractions of 1.5 ml from the top and combined the fractions corresponding to the 20S-40S region.

### Purification of LtPCC by streptavidin affinity pulldown

We supplemented glycerol gradient fractions with octylglucoside to 2 mM. On a nutating mixer, we incubated the fractions on Strep-Tactin®XT magnetic beads in a Binding Buffer (50 mM Tris-HCl, pH 8.0, 2 mM OG, 1 mM EDTA, 100 mM KCl) at 4°C for 1 hour. We washed the beads twice, with 5 ml of Binding Buffer each time. Then, we incubated the beads for 10 minutes and at 4°C, in 0.2 ml of Elution Buffer (20 mM Tris-HCl, pH 8.0, 100 mM biotin, 1 mM EDTA, 100 mM KCl, 2 mM OG). Using Zeba^™^ Spin Desalting Columns, 7K MWCO (0.5 ml), we exchanged the 130 μL of purified material into the Sample Buffer (20 mM Tris-HCl, pH 7.5, 60 mM KCl, 5 mM MgCl2, 1 mM DTT, 5 mM OG). We centrifuged the sample for 10 minutes at 21,000g. We stored the supernatant on ice before cryoEM grid preparation.

### CryoEM sample preparation and image acquisition

We first used PELCO easiGlow^™^, with a target current of 15 mA, to glow discharge Lacey carbon cryoEM grids with a 2 nm continuous carbon film (Ted Pella) for 45 seconds. Then, we applied 2.5 μL of sample onto the grids. We waited for 5 seconds before blotting the grids for 4 seconds with blot force 0, at 100% humidity and 4°C. After we blotted the grids, we used a FEI Mark IV Vitrobot (Thermo Fisher Scientific) to plunge-freeze the grids into liquid ethane. We stored the grids in a liquid nitrogen dewar until cryoEM image acquisition.

We loaded and imaged the cryoEM grids through a Titan Krios (Thermo Fisher Scientific) with a Gatan Imaging Filter Quantum LS and K3 camera, operated at 300 kV. We recorded movies at a pixel size of 0.55□Å/pixel with SerialEM [41], by electron counting in super-resolution mode. We set an exposure time of 2 seconds, fractionated to 40 frames, and a defocus range between −2.5 to −1.5 μm. We had an estimated total dosage of 40 e-/Å^2^. We collected 3,328 movies.

### CryoEM image processing

#### Reconstruction of cryoEM maps

To produce drift-corrected and dose-weighted micrographs, we processed the movies with MotionCor2 [42]. After motion correction, the movies had a calibrated pixel size of 1.1 □Å. Due to severe drift of the first frame, it was discarded. We determined the defocus values of the micrographs with Gctf [43]. We first used Warp’s BoxNet [44] to automatically pick particles. Afterwards, in RELION 3.1 [45], we performed reference-free 2D classification, and selected the classes with features for 3D classification with D3 symmetry. After 3D classification, we selected good classes, with a total of 8214 particles, to train a topaz model. Then using this model, we picked particles through topaz. We used RELION 4.0 [46] to perform multiple rounds of reference-free 2D classification of the topaz-picked particles. From the 2D classes, we selected 15859 particles for 3D refinement in RELION 4.0 [46]. We used a map from the previous 3D classification of BoxNet particles as the reference map. After 3D refinement with D3 symmetry, we obtained a map of 3.49 Å. We iteratively refined the 3.49 Å map through CTF refinement and 3D refinement [46]. After all processing, its final resolution was 3.2 Å (Figure S4).

We then performed D3 symmetry expansion on the particles from D3 3D refinement. Using a mask of one α-subunit, we performed 3D classification with C1 symmetry. For 3D classification, we skipped both angular and offset search in RELION 4.0 [46]. We then performed 3D refinement with C1 symmetry and local search, obtaining a α_5_β_6_ map of 10.6 Å after postprocessing (Figure S4). Afterwards, we performed C3 symmetry expansion on the particles from D3 3D refinement. Using a mask of two α-subunits, we performed 3D classification with C1 symmetry, skipping both angular and offset search [46]. We then performed 3D refinement with C1 symmetry and local searches, obtaining a α_4_β_6_ map of 18.4 Å after postprocessing (Figure S4).

#### Sorting method

The sorting method is based on the focused classification and symmetry expansion method described here [47]. We performed 3D reconstruction with D3 symmetry using all the particles to obtain the *rot, psi* and *tilt* Euler angles of each particle. We expanded the particles for D3 symmetry and masked out one α-subunit from the D3 symmetry 3D reconstruction. We then performed focused classification without angular and offset search on the symmetry expanded particles, with two classes. We visually inspected which class was empty. We counted how many symmetry-expanded particles were in the empty class, which corresponds to the number of unoccupied α-β binding sites in the PCC particle (Table 1). From this, we calculated the number of LtPCC particles sharing an oligomeric formula.

#### Atomic modeling and model analysis

We first modeled and refined one α-subunit and one β-subunit in Coot [48], based on the AlphaFold [32] prediction for LtPCC. Then, through ChimeraX [49], we duplicated the subunits to D3 symmetry and fit them into the LtPCC cryoEM map reconstructed with D3 symmetry. The model was then refined first through Phenix’s real space refine function [50], and manually checked in Coot [48]. This model was fitted into the α_5_β_6_ and α_4_β_6_ maps. Based on the two maps, one and two α-subunits were removed from the original model, respectively. Hydrogen bonds and interfacial residues were analyzed through ChimeraX [49] and verified in PISA [51]. The binding affinity was calculated through the PRODIGY web server [52,53]. The sequence alignment between LtPCC and HsPCC was done through Clustal Omega [54] and visualized through ESPript 3 [55].

## Data and code availability

The α_6_β_6_, α_5_β_6_ and α_4_β_6_ cryoEM maps have been deposited in the Electron Microscopy Data Bank under accession numbers EMD-XXXX, EMD-XXXX, and EMD-XXXX, respectively. The coordinates of LtPCC models have been deposited in the Protein Data Bank under accession number XXXX. All aforementioned deposited data are publicly available as of the date of publication. This paper does not report original code. Any additional information required to reanalyze the data reported in this paper is available from the lead contact upon request.

## Acknowledgments

Our research was supported in part by grants from the U.S. National Institutes of Health (R01GM071940 to Z.H.Z. and R01AI101057 to R.A.). We acknowledge the use of resources at the Electron Imaging Center for Nanomachines of UCLA supported by U.S. NIH (S10RR23057 and S10OD018111) and U.S. NSF (DMR-1548924 and DBI-133813). We thank William Lan for initial efforts in model building.

## Author contributions

Z.H.Z. and R.A. initiated and supervised the project. I.A. prepared the sample. Y-T.L. carried out cryoEM imaging. Y-T.L. and J.K.J.L. performed data processing. J.K.J.L., Y-T.L. and J.J.H. analyzed the data, made illustrations, and wrote the original draft. Z.H.Z., Y-T.L., J.K.J.L. and J.J.H. finalized the manuscript. All authors reviewed and approved the paper.

## Declaration of interests

The authors declare no competing interests.

---

**Figure S1.**
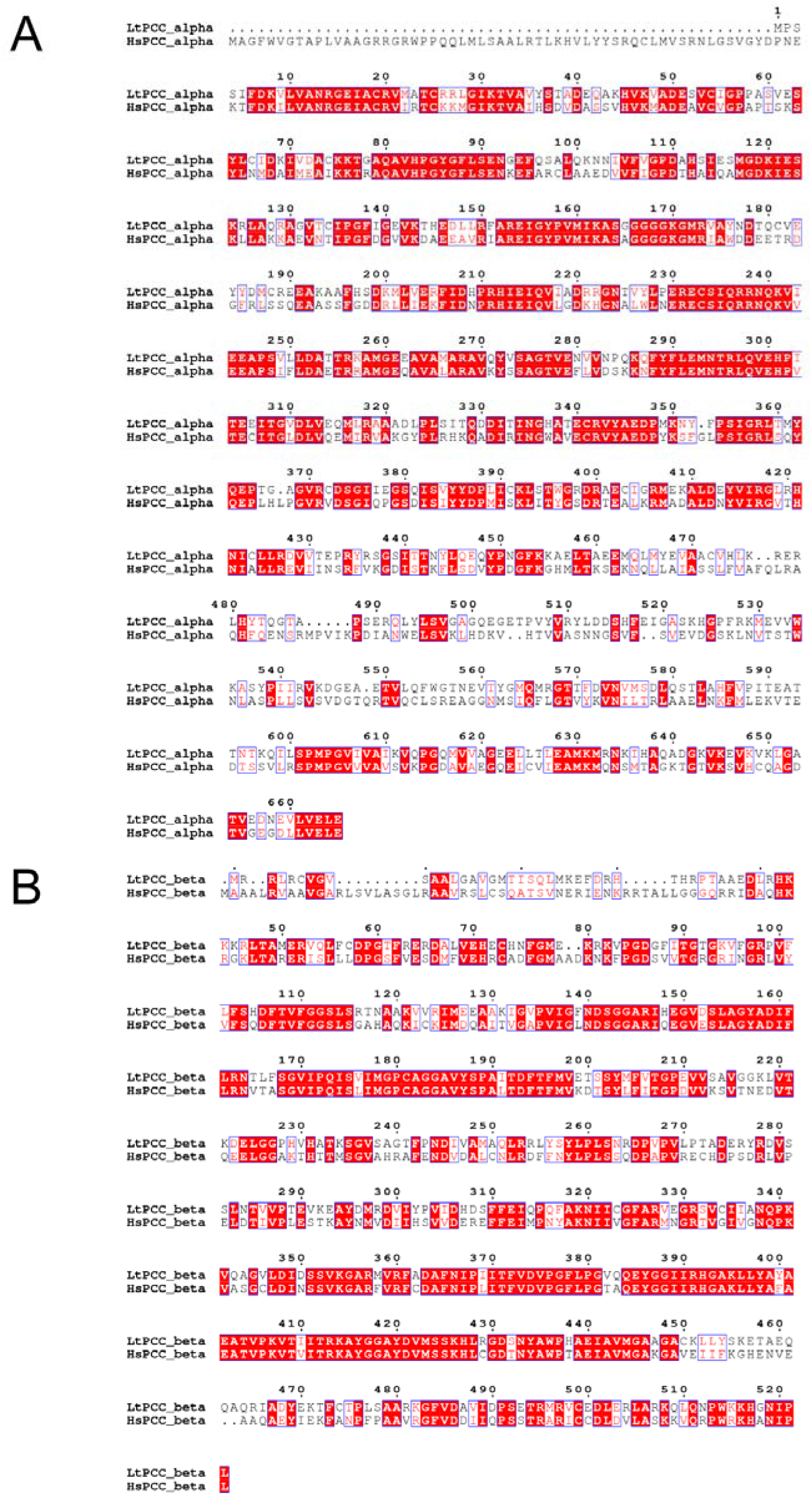
Sequence alignment of LtPCC α-subunits (A) and β-subunits (B) with HsPCC α-subunits and β-subunits. There is a dot on top of every ten residues. Blue-outlined boxes denote similar residues. Red-background denotes identical residues.

**Figure S2.**
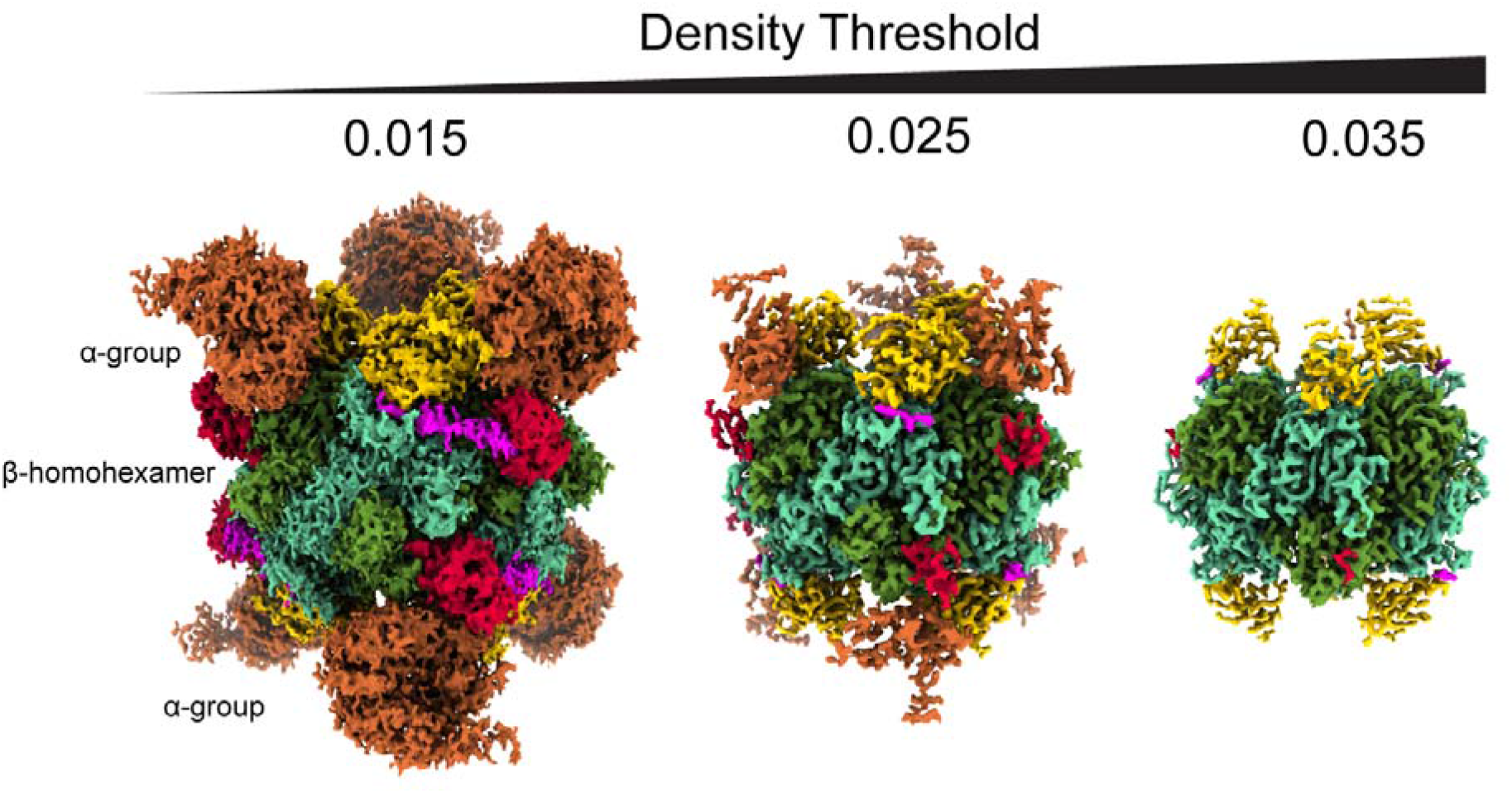
In D3 reconstruction, LtPCC α-subunits exhibit flexibility and/or lower occupancy at high threshold while the β-homohexamer remains stable.

**Figure S3.**
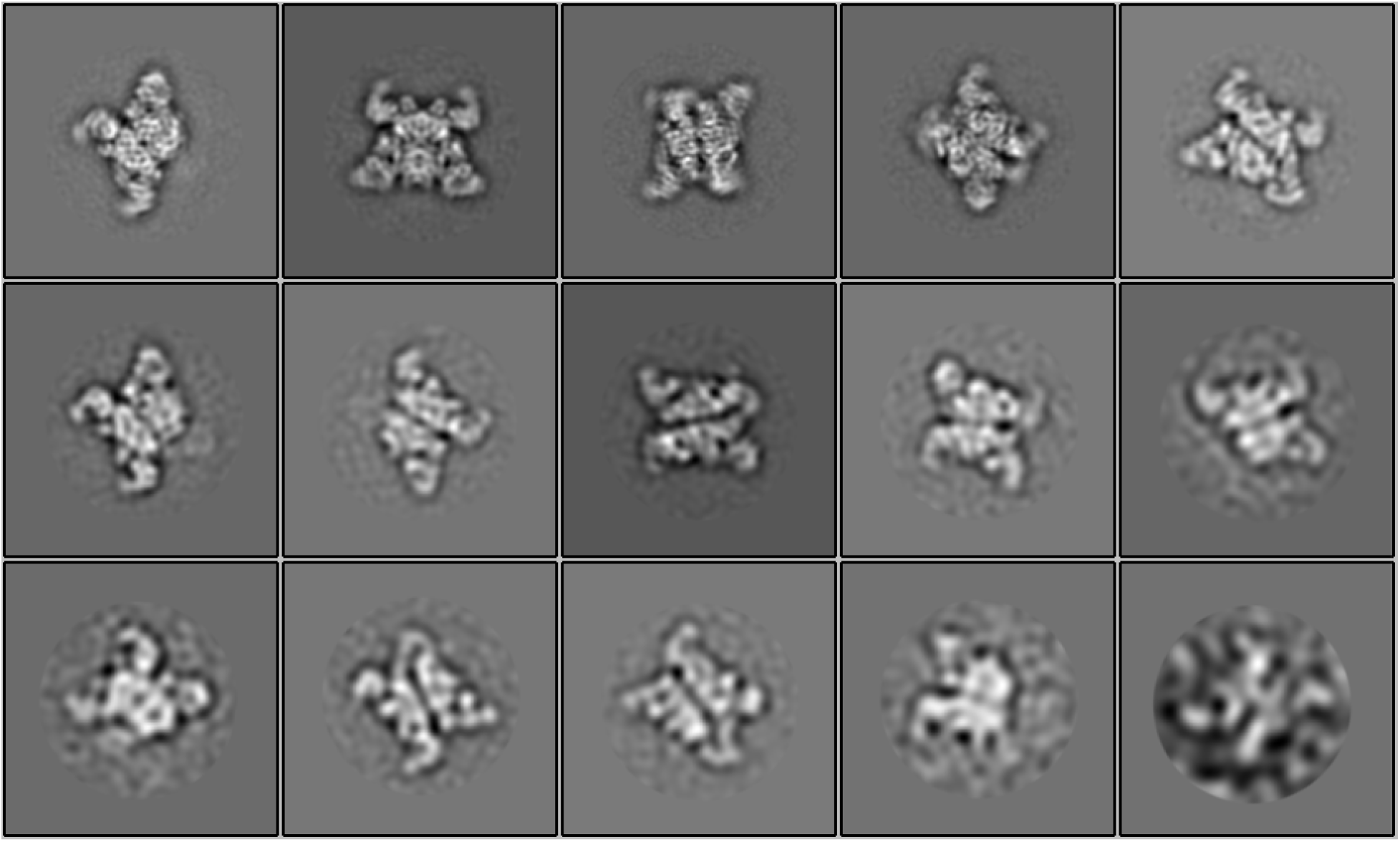
All selected 2D classes for LtPCC reconstruction with D3 symmetry.

**Figure S4.**
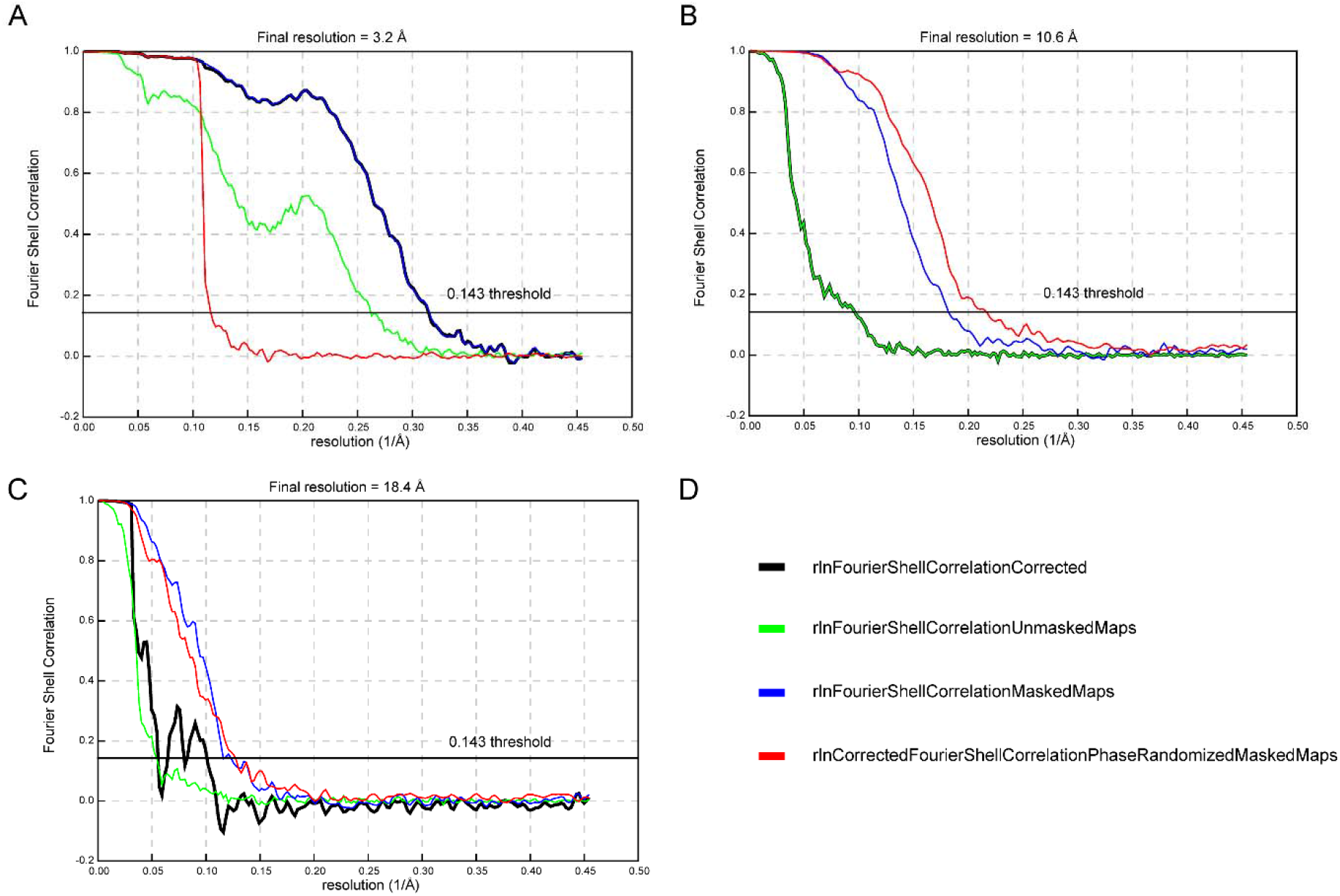
Estimation of resolution for the three cryoEM maps. Fourier shell correlation curves as colored in (D) for the α_6_β_6_ map (A) α_5_β_6_ map (B) α_4_β_6_ map (C).

## Notes

### Competing Interest Statement

The authors have declared no competing interest.

